# A comprehensive analysis of grazing: Improving management for environmental health

**DOI:** 10.1101/2023.06.06.543944

**Authors:** Talia E. Niederman, Ashley L. Xu, Lindsay M. Dreiss

## Abstract

In an effort to combat the dual climate and biodiversity crises, the international community has put forth targets to reduce emissions and protect species. Habitat degradation is at the fore in driving species extinctions and carbon loss, creating an urgent need to examine our land-use practices if we are to meet international goals. Individual nations will have differing needs and approaches to reaching these objectives based on their landscapes, industries, and levels of historic overuse. In the United States, grazing is the leading land-use, making up approximately one-third of the country. Here we present a broad-scale analysis of how grazing effects the environment and examine how this land-use plays out across the U.S. This review can help policy makers and land managers strategically assess grazing lands as we work towards a national landscape that supports equitable ecosystem services, sustainable livelihoods, and climate resilience.

**Highlights:**

- Livestock grazing can have a multitude of effects on local ecosystems often causing habitat degradation and loss. As this land-use makes up a third of U.S. lands, improving grazing practices could provide significant benefits for the environment.
- To our knowledge, this is the largest review of U.S. grazing to date. We explore how grazing affects six elements of the environment across the country.
- While the majority of literature discusses negative implications related to grazing, our results indicate that regions are affected differently, and that management and livestock-type also contribute to the overall environmental impact.

## Intro

On our current course, we are at risk of losing 1 million species to extinction (IPBES 2019) and reaching temperatures 3.2°C above preindustrial levels by the end of the century (IPCC 2023). In the midst of these dual and compounding crises, nature-based solutions will be essential to shifting our trajectory (Pörtner *et al*. 2023). It is therefore imperative we understand how our use of lands and waters affects ecosystems. Within the United States, grazing is the most extensive land-use, covering roughly one-third of the country (Bigelow and Borchers 2017). Recognizing how grazing influences environmental processes can help inform management for this widespread practice. Establishing a responsible U.S. grazing regime is essential to supporting healthy biodiversity and ecosystem services as well as human livelihoods and local economies.

While there is a wealth of literature on grazing, many reviews take a regional focus (e.g., Stahlheber and D’Antonio 2013) or a global one (e.g., Schieltz and Rubenstein 2016).

Understanding grazing trends at the national level is essential for salient policy and land management. To address this, we conducted an extensive review of literature discussing grazing across U.S. ecosystems from the past two decades, which to our knowledge, is the most comprehensive review of U.S. grazing to date. This synthesis can demonstrate how grazing affects key environmental processes and help improve management practices.

This review specifically focused on six environmental factors: water quality, erosion, nutrient cycling, fire regime, wildlife and habitat, and soil carbon levels. Through a synthesis of the literature, we report on how grazing effects these factors, define relationships between grazing and each factor by ecoregion, and analyze the influence of livestock-type. This review is particularly timely as national efforts to conserve 30% of lands and waters by 2030 progress and conversations often diverge on what kinds of land-use should count toward conservation goals.

Understanding how grazing affects ecosystems across the U.S. can support policy makers in formulating conservation strategies and identifying which lands should qualify for 30×30. So too, as the Bureau of Land Management (BLM) moves to prioritize conservation at a higher level, comprehensive grazing information is essential as a major use of BLM lands (BLM 2023). Locally, this overview can guide land managers and conservation practitioners to ask further questions about their regions of focus and place them within a broader landscape framework.

## Methods

We conducted a systematic literature review to examine how grazing in the U.S. effects local environments. Search terms encompassed six environmental factors identified through consultation with experts: water quality, erosion, nutrient cycling, fire regimes, wildlife and habitat, and soil carbon (see supplemental material, Table S1). The search was limited to literature regarding North American ecosystems with data relevant to grazing in the U.S. published from 1999-2021. We excluded papers solely discussing livestock greenhouse gas emissions as these speak to broader environmental degradation through climate change; this review focused on the more site-specific effects of grazing (e.g., from the six factors identified above).

We used Scopus as our primary search engine supplemented with results from Google Scholar and contributions from experts. For each search (Table S1), we screened the first 1,000 results (relevant articles were depleted well before this point). We then removed duplicates, irrelevant papers, and those with completely inconclusive results leaving 726 papers.

For our qualitative analysis, we recorded whether papers relayed “positive,” “dependent/positive,” “dependent,” “dependent/negative,” “negative,” or “neutral” effects of grazing on the environment. “Positive” and “negative” were assigned to papers who fell most neatly into these categories. “Dependent” encompassed papers where grazing might have different outcomes depending on the situation. Papers discussing varying extents of positive or negative effects due to circumstance were labeled “dependent/positive” or “dependent/negative”. When grazing did not have an effect, papers were categorized as “neutral” (examples in Table S2). A paper discussing a positive or negative element of grazing doesn’t necessarily imply grazing is wholly positive or negative, but the aggregate of results portrays broader trends.

We categorized overall results as well as for the six environmental factors (as relevant) and recorded year published, livestock, species, and ecoregions discussed (EPA Level I). We did not record ecoregion from large-scale reviews or studies discussing wide areas with unclear boundaries. When a study site’s ecoregion wasn’t completely apparent, we used context with our best judgment.

We analyzed the overall results and trends for the six environmental factors. We also looked at trends by ecoregion (ecoregions with >25 results), environmental factor for each ecoregion (environmental/ecoregion with >5 results), species effected, and livestock. To understand how environmental factors play out by ecoregion we calculated representative values for each intersection based on an average of results (more details in Table S3).

## Results

Trends for grazing varied by ecoregion, livestock type, species, practice, and management objective. Overall (N=726), 46% of papers described negative-leaning trends (33% “negative”, 13% “dependent/negative”), 26% were dependent, 20% positive-leaning (13% “positive”, 7% “dependent/positive”), and 9% neutral. The breakdown of results did not appear to change meaningfully over time. There was more negative-leaning literature for all environmental factors, but the highest concentrations were for water quality (84%) and erosion (78%). The highest concentrations of positive-leaning papers were for fire regime (38%), habitat (21%) and soil carbon (21%; Figure 1).

**Figure 1:**
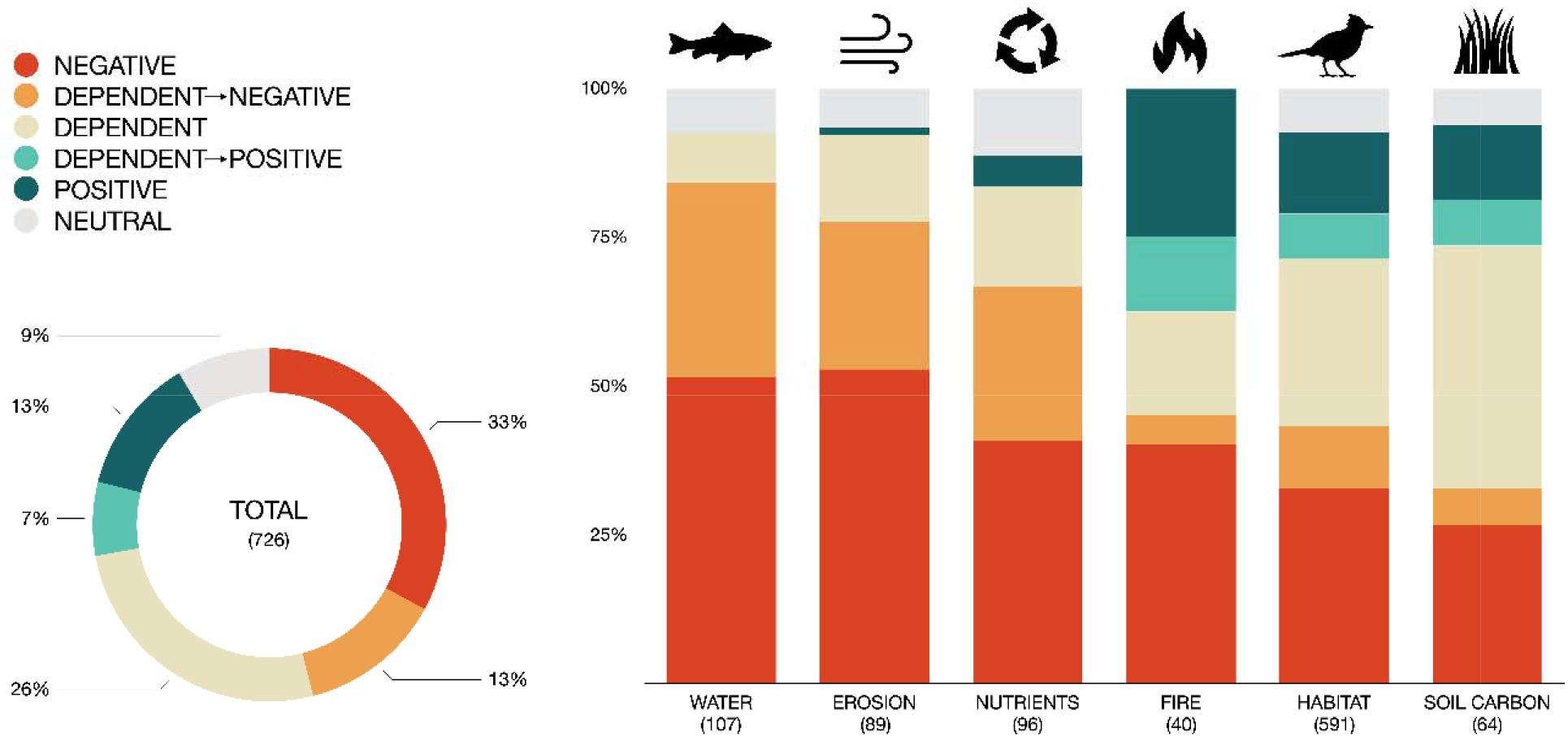
Grazing papers were classified on a continuum of negative to positive based on the effects they reported. Here we show the breakdown of results for grazing overall and specific to six environmental factors (water quality, erosion, nutrient cycling, fire regime, wildlife and habitat, and soil carbon levels). Factors are listed from left to right in terms of highest percentage of negative-leaning papers (“negative” and “dependent/negative”). Sample size is included in parentheses.

All ecoregions had more papers reporting negative impacts than positive, except for the Great Plains which was equal (30% each). The following ecoregions had a higher percentage of negative-leaning papers than the overall average: Northwestern Forested Mountains (62%), Southern Semi-arid Highlands (60%), North American Deserts (57%), Temperate Sierras (53%), and Eastern Temperate Forests (53%). While no ecoregions had majority positive-leaning papers, the following ecoregions had a greater percentage of positive-leaning papers than the overall average: the Great Plains (30%) and Southern Mediterranean California (30%).

Looking at environmental factor by ecoregion, nutrient cycling and carbon storage in the North American Deserts had the highest percentage of negative-leaning papers. The highest percentages of positive-leaning papers were for fire regime and carbon storage in the Great Plains (Figure 2b).

**Figure 2:**
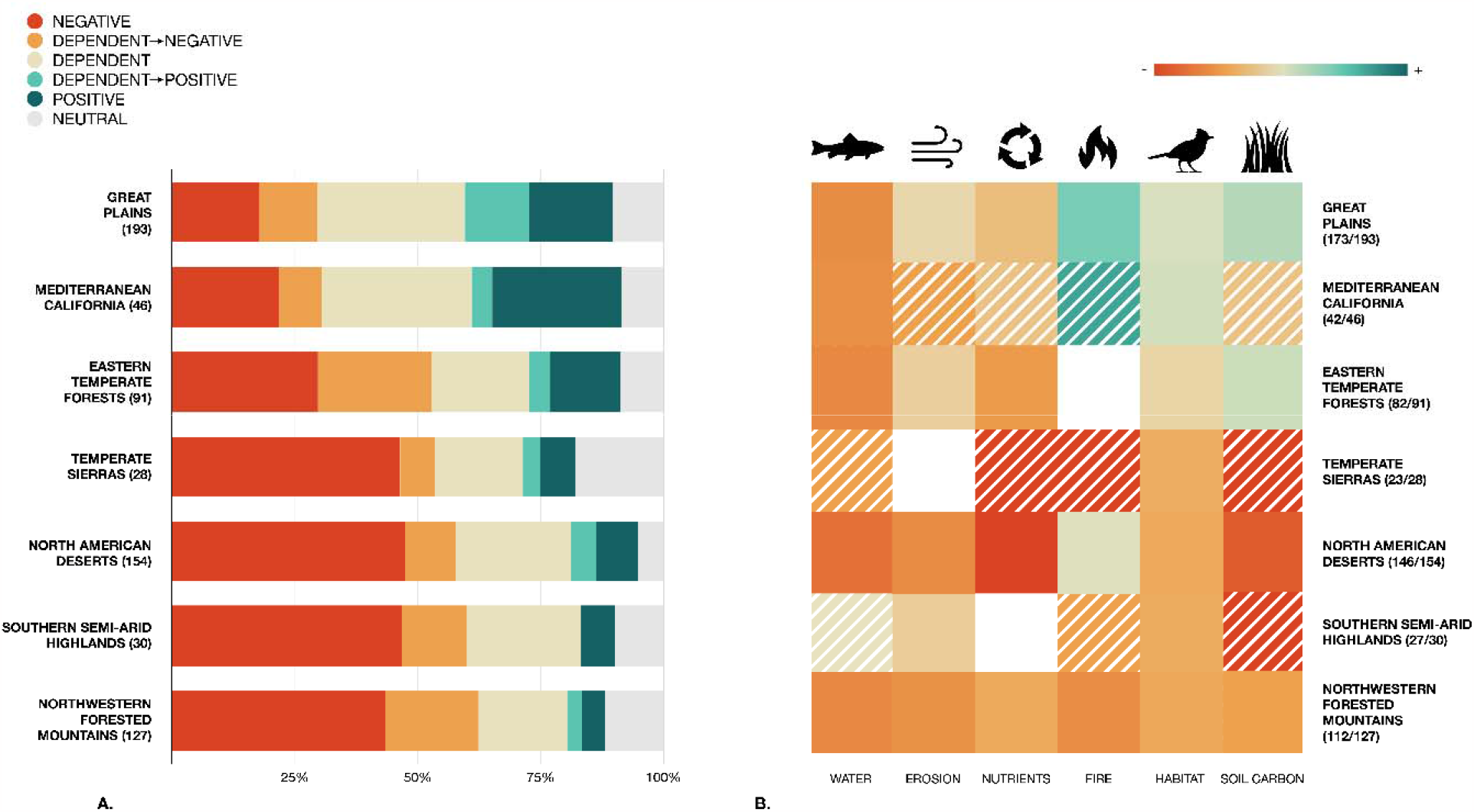
On the left (2a) grazing trends in the literature are broken down by ecoregion (for ecoregions with >25 results). The results are listed from top to bottom in terms of lowest to highest percentage of negative-leaning papers (“negative” and “dependent/negative”). Sample size is included in parentheses. To the right (2b), ecoregion trends are broken down further by the six environmental factors (white boxes indicate no results and hatched boxes indicated an environmental factor/ecoregion crossover with <5 results). Some these subdivisions were much more robust than others see supplementary materials (Figure S1) to view these results with the added dimension of sample size.

Of livestock, literature was most likely to describe a positive effect for bison grazing (73% positive-leaning and 7% negative-leaning). Other livestock (cattle, sheep and goats, and other groupings) were more comparable to each other (positive-leaning: 19%, 22%, and 14% respectively and negative-leaning: 47%, 53%, and 48% respectively; Figure 3). Literature on bison grazing was generally restricted to the Great Plains. Within Great Plains literature, bison still had a drastically larger percentage of positive-leaning papers (69% positive-leaning, 9% negative-leaning; Figure 3) compared to papers on grazing of other livestock species (25% positive-leaning, 32% negative-leaning).

**Figure 3:**
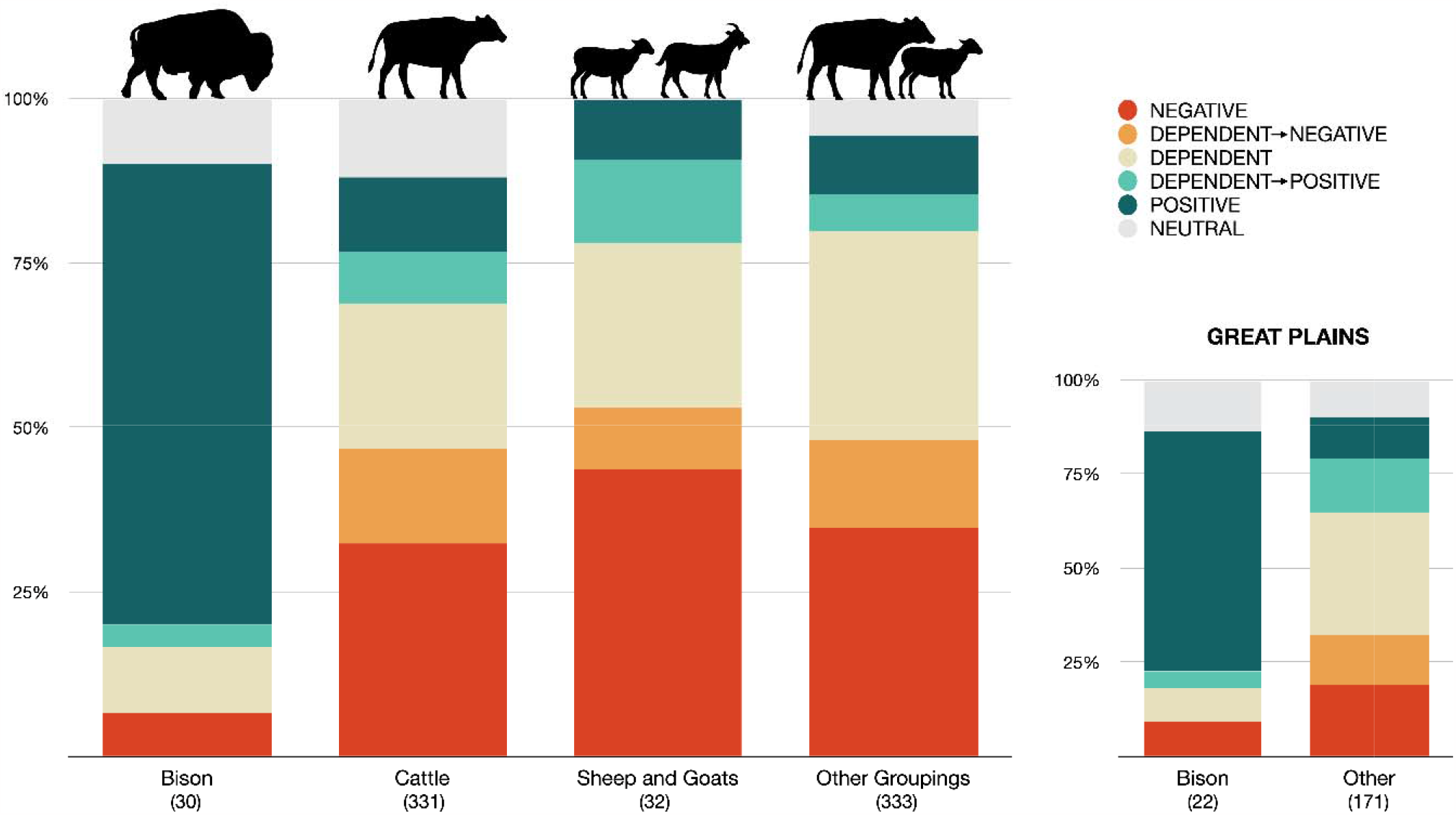
Here we show grazing trends broken down by livestock-type. Bison had the greatest percentage of positive papers. Bison were mainly restricted to the Great Plains, so we also show the comparison of bison to other livestock specifically for this ecoregion. Cattle had by far the most papers (including the cattle category as well as many others that fell into the “other groupings” category) so this should be noted in interpreting results. Sample size is included in parentheses.

Plants and wildlife showed similar trends to each other, but there was a slightly higher percentage of papers describing negative effects for vertebrates (44%) than invertebrates (32%). Of vertebrates, fish had the highest percentage of negative-leaning papers (74%), followed by mammals (56%), amphibians (50%), birds (34%), and reptiles (31%). Riparian and wetland species seemed to be most negatively affected plants (74%; Figure 4). We urge caution in putting too much stake in these species-specific results as they encompass broad groups and effects can vary widely per species.

**Figure 4:**
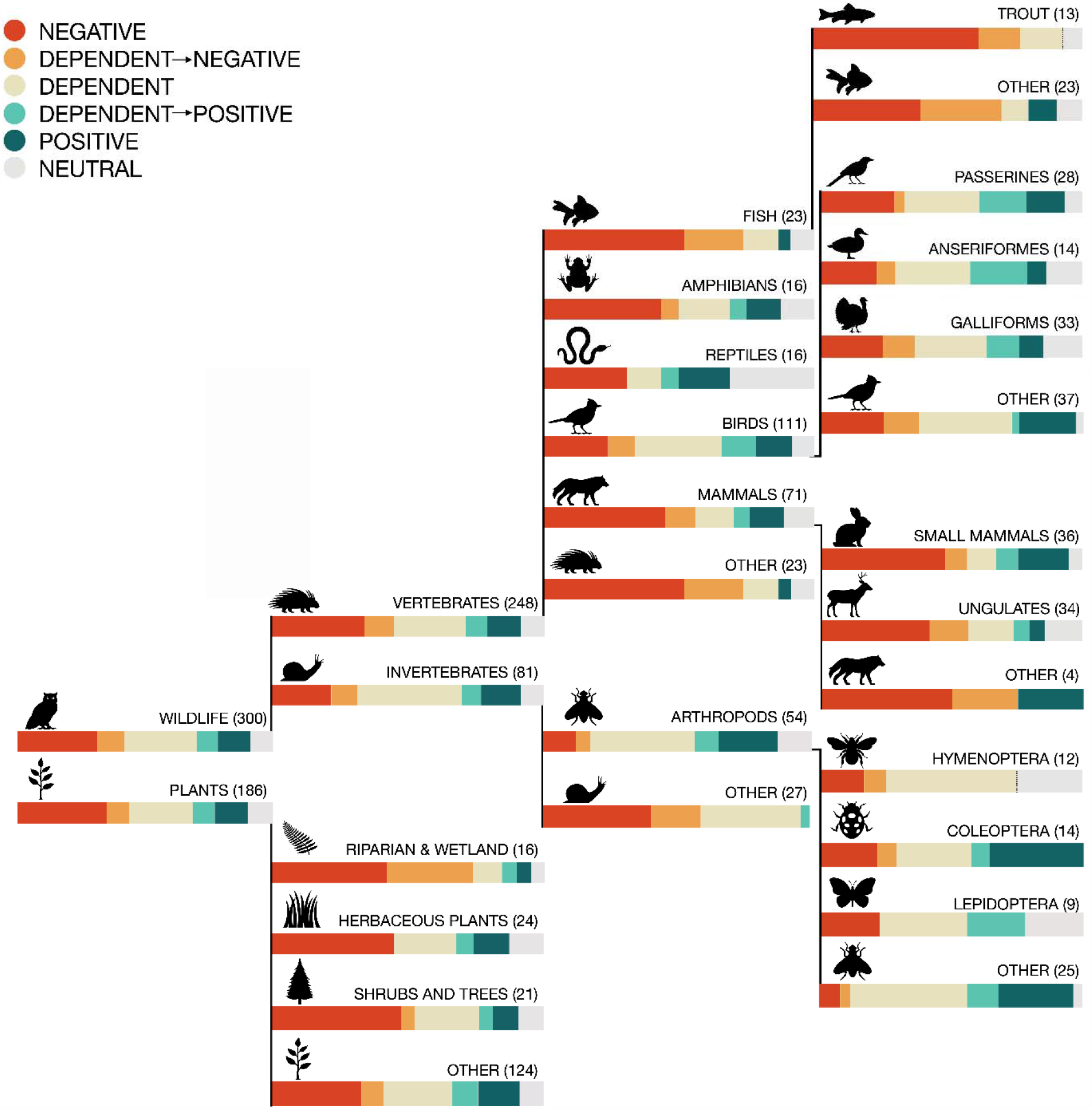
This tree shows a breakdown of grazing trend by species grouping. This is just a sample of species that may be affected by grazing and these results should simply serve to speculate on broad trends regarding these groups as effects on individual species can vary widely. Sample size is included in parentheses.

## Discussion

In this review of grazing within U.S. ecosystems, papers discussing negative trends more than doubled the positive. Negative-leaning papers also outnumbered dependent and neutral papers (although authors may be less likely to publish their work if they found no significant effect, thereby decreasing the number of neutral papers). These results indicate that while grazing may benefit the environment in some instances, most of it currently does not. This is likely due to a combination of environmental incompatibility (e.g., Fick *et al*. 2020) and poor management (e.g., Krall *et al*. 2021). Demonstrating this, only 34% of grazed BLM lands meet rangeland health standards or are making significant progress towards them (BLM 2019). Fortunately, the negative effects of grazing can be alleviated at times. Understanding situations in which grazing is detrimental or beneficial could help inform both local and national land management decisions in the U.S. Here we provide some insight into how grazing affects different environmental factors. Although we discuss these separately, they are often interconnected.

### Water Quality

Papers addressing grazing impacts to water quality widely reported negative outcomes (84%). Grazing can elevate water temperatures, increase sedimentation and turbidity, introduce harmful pathogens into water bodies, and cause eutrophication (Schindler *et al*. 2012; O’Callaghan *et al*. 2019; Kovach *et al*. 2019). How severely water bodies are affected depends on factors like climate, stream size, livestock access, grazing intensity, soil drainage, and ambient water quality (O’Callaghan *et al*. 2019). Cattle seek out water more frequently during hot weather, which can intensify water quality issues during these times; they also preferentially defecate in water bodies, contributing to degradation (Pandey *et al*. 2009; O’Callaghan *et al*. 2019). Comparatively, bison are more heat-tolerant and spend less time in water (Grudzinski *et al*. 2018).

Diminished water quality from grazing is a widespread issue that is important to address as clean water is essential to both people and the environment. Strict management may be particularly necessary in areas with imperiled aquatic species (Clark Barkalow and Bonar 2015). Livestock exclusion from riparian areas and well-placed off-stream water troughs can help keep livestock away from fresh water bodies ameliorating water quality (Johnson *et al*. 2016; Krall *et al*. 2021). As climate change exacerbates, replacing cattle with bison may be strategic in appropriate ecoregions as bison have higher heat tolerance making them less likely to wallow (Grudzinski *et al*. 2018).

### Erosion

Literature reporting grazing’s impact on erosion also had a high percentage of negative-leaning papers (78%). Grazing can increase bare ground prompting soil compaction and erosion (Centeri 2022). While grazing causes erosion in all ecoregions, arid regions seem to be some of the worst affected (Fick *et al*. 2020); these effects are exacerbated by drought (Nauman *et al*. 2018), which will become more prevalent with climate change. Grazing-induced erosion and stream bank degradation is also an issue for riparian habitats and can contribute to water quality issues (Zaimes *et al*. 2021).

Proven ways to prevent or lessen erosion include implementing rotational over continuous grazing, lowering stocking densities, and resting erosion-prone areas after disturbance (Lyons *et al*. 2000; Tufekcioglu *et al*. 2013; Clark *et al*. 2018). In riparian ecosystems, excluding grazing or avoiding the main growing season may help improve bank stability (Dalldorf *et al*. 2013) as well as practices for improving water quality mentioned above. Arid areas with high susceptibility to erosion should likely be avoided for grazing.

### Nutrient Cycling

Most papers discussing livestock’s effect on nutrient cycling described negative outcomes (67%). Livestock excrete high levels of nitrogen and phosphorous, which can contribute to water quality issues and eutrophication (Schindler *et al*. 2012). In nutrient-poor areas, livestock may have a particularly negative effect on nutrient cycles causing nutrient loss and reduced soil fertility (e.g., North American Deserts Fernandez *et al*. 2008). Grazing can also facilitate broader-scale nutrient disruptions (e.g., increased eolian dust deposition across ecosystems; Neff *et al*. 2008).

Stocking rate and grazing intensity largely influence nutrient runoff with lighter-grazed areas hosting more vegetation and less runoff (Tufekcioglu *et al*. 2013; Burt *et al*. 2013). Some other management tools suggested included riparian buffers, lower grazing elevation, and improved livestock nutrition (Derlet *et al*. 2012; Beck and Gregorini 2021; Dunn *et al*. 2022).

### Fire Regime

Depending on ecoregion and management, grazing may have positive or negative effects on fire regime. It should be noted that only papers discussing grazing’s effect on fire were included in this section of the analysis and that some papers discussing the effects of fire coupled with grazing as a management tool (pyric herbivory) may only be included in the “wildlife and habitat” category if grazing didn’t directly affect fire regime. Nevertheless, we generally discuss pyric herbivory in this section to communicate management recommendations.

In some cases, grazing can help prevent extreme fires by regulating plant growth or creating fuel breaks (Huntsinger and Barry 2021). Pyric herbivory can benefit ecosystems that have evolved with it and promote a heterogeneous landscape beneficial to some species (Wilcox *et al*. 2022). However, improperly managed grazing may over-suppress fires resulting in habitat degradation; overgrazing and fire suppression of the western U.S. during Euro-American colonization notoriously led to mismanaged forests resulting in overgrown understories and catastrophic fires (Addington *et al*. 2018). The literature was mixed on whether grazing increases the prevalence of invasive species elevating extreme fire risk (e.g., Williamson *et al*. 2020) or controls invasives lowering it (e.g., Huntsinger and Barry 2021). This may be dependent on the invasive species and its level of establishment.

While several ecoregions had insufficient data to see trends of grazing on wildfire regime, the Great Plains seemed the most likely to see positive outcomes from pyric herbivory. Historically, the Great Plains were shaped by fire and bison so well-managed pyric herbivory may be beneficial for this ecosystem (Fuhlendorf *et al*. 2009). In cases where it could be beneficial, pyric herbivory should be applied strategically so as not to harm habitat or sensitive species (Davies *et al*. 2016). Site-specific factors like recent disturbance and landscape matrix should also be taken into account (Beschta *et al*. 2004; Fuhlendorf *et al*. 2009).

### Wildlife and habitat

Species and habitats may be affected by grazing in myriad ways (Figures 3 & 4). Our results showed that aquatic habitats and more arid ecosystems are particularly vulnerable. Grazing can threaten habitat through numerous mechanisms including, but not limited to, degraded biocrusts, altered plant composition, increased desertification, and other ecosystem changes discussed above (Clements 2004; Chambers *et al*. 2017; Root *et al*. 2020). Overall, areas that evolved without large native grazers may be less suitable for grazing (Bakker *et al*. 2006). Grazing may also threaten individual species through habitat degradation or in more targeted ways [e.g., disease transmission (Cassirer *et al*. 2018), human-wildlife conflict, or fencing (Jakes *et al*. 2018)]. It can also compound upon other threats to species (Cisneros-Pineda 2020). Invasive species are a major threat to wildlife and habitat, but our review found mixed results on grazing and invasives. Overall, there were more papers discussing negative effects of livestock in regards to invasive species (e.g., Kauffman *et al*. 2022), but there were also positive cases (e.g., Dornbusch *et al*. 2020) indicating that the effect of livestock on invasives may be situational.

Alternatively, some habitats and species may benefit from well-managed grazing such as within the Great Plains or Mediterranean California (Gennet *et al*. 2017; Wilcox *et al*. 2022). For these ecosystems, grazing promoting heterogeneity may support certain species such as grassland birds (Gennet *et al*. 2017). Results indicated particularly positive trends for areas grazed by bison on the Great Plains. While other species of livestock also had fewer negative effects on this ecoregion, bison were by far the most likely to have positive outcomes.

Grazing can have wide-ranging effects on ecosystems so choosing appropriate grazing areas and employing suitable management methods, such as rotational grazing, can significantly influence the effects of this land use (Hillenbrand *et al*. 2019). Bison may have fewer negative effects than cattle (e.g., less riparian degradation) so managing with bison could produce better environmental outcomes in some situations (Grudzinski *et al*. 2018). Evaluating land on a case-by-case basis to determine grazing feasibility and management practices is essential.

### Soil carbon

Our results indicated that the effects of grazing on soil carbon is extremely variable. Grazing seems most likely to promote Great Plains soil carbon stores (e.g., Bork *et al*. 2020) whereas it may be most harmful for North American Desert stores (e.g., Fernandez *et al*. 2008). Grazing intensity can influence whether carbon is lost or stored (Ma *et al*. 2021) and continuous grazing may be harmful for carbon storage objectives (Byrnes *et al*. 2018). In situations where grazing could enhance carbon storage, this should be weighed against GHG emissions by livestock to determine net gain or loss of carbon.

## Conclusions

The existing body of literature on grazing in the U.S. demonstrates that it is having consequences for ecosystem health. Although grazing often has negative repercussions, there are instances in which it may provide benefits. Ultimately, the environmental outcome is likely influenced by both ecosystem characteristics and management practices While this review illustrates broad trends, it is important to assess land management decisions on a local basis with a thorough understanding of imperiled species and ecosystem needs.

It is essential to note that this review focuses on scientific literature and may fail to include critical perspectives. In particular, American Indians have lived on this land since time immemorial and many Tribes have important insight into grazing practice, particularly regarding bison. Across the U.S., vegetative regimes have been largely altered through the loss of Indigenous management (Huntsinger and Barry 2021). While we included search terms surrounding Traditional Ecological Knowledge (TEK) and Indigenous grazing practices (Table S1), we believe that this review lacks the guidance of these voices due to the general dearth of representation in scientific literature. Tribes are currently leading important work in sustainable grazing such as through the Buffalo Treaty, an intertribal alliance that aims to promote bison across large swathes of North America (Blackfeet Nation *et al*. 2014). We hope that this review can be used in conjunction with other forms of knowledge, including TEK, to gain a comprehensive understanding of the impacts of grazing throughout the U.S.

In forming broad-scale conservation policies and targets, it is key to have a comprehensive understanding of grazing across the U.S. As the most prevalent U.S. land-use and a major presence on public lands, accounting for the environmental effects of grazing across the country is essential to well-founded decision making. Practitioners should consider whether an ecosystem is appropriate for grazing and if so, how to graze in a way that promotes conservation values and limits degradation. While there are some instances in which grazing may enhance certain habitat objectives, it is vital to recognize the many environmental threats associated with grazing, and grazed lands should not be assumed to qualify for biodiversity goals. Through meaningfully revisiting and reshaping grazing within the U.S., we can make strides towards improved land management as we aim to protect biodiversity and improve the sustainability of human livelihoods.

## Acknowledgments

We thank S. Ozbenian, J. Malcom, S. Cantrell, and P. Nelson for assistance formulating project goals and methodology.

We would also like to acknowledge that recent work in the field of conservation science and STEM at large has identified a bias in citation practices such that papers from women and minority groups are relatively under-cited (see Larivière et al. 2013, Rudd et al. 2021 and others). As we did not proactively choose references that reflect the diversity of the field, we recognize the biases that may have been unintentionally introduced. We look forward to future work that can help us to better understand how to support equitable practices in conservation science.

## Conflict of interest

The authors declare no conflicts of interest.

## Data Availability Statement

All data used for this study can be found at: https://osf.io/msrtg/

## Notes

### Competing Interest Statement

The authors have declared no competing interest.

