## supplemental material for "A comprehensive analysis of grazing: Improving management for environmental health"

**SUPPLEMENTAL MATERIALS**

**Table S1.** Search terms for our systematic literature review of North American grazing regimes.

| Species |  | Conservation |  | Grazing |  | Location |
| --- | --- | --- | --- | --- | --- | --- |
| Livestock |  | Biodiversity |  | Grazing |  | North America |
| *OR* |  | *OR* |  | *OR* |  | *OR* |
| Cattle | AND | Wildlife | AND | Graze | AND | United States |
| *OR* |  | *OR* |  | *OR* |  | *OR* |
| Cow |  | Ecosystem |  | Agriculture |  | U.S. |
| *OR* |  | *OR* |  | *OR* |  | *OR* |
| Beef |  | Habitat­­ |  | Natural grazing regime |  | US |
| *OR* |  | ________________ |  | *OR* |  | *OR* |
| Bison |  | Wildfire |  | Prescribed grazing |  | Mexico |
| *OR* |  | *OR* |  | *OR* |  | *OR* |
| Sheep |  | Fire­ |  | Regenerative grazing |  | Canada |
| *OR* |  | ________________ |  | *OR* |  | *OR* |
| Horse |  | Soil carbon |  | Regenerative agriculture |  | Montana |
| *OR* |  | *OR* |  | *OR* |  | *OR* |
| Alpaca |  | Soil |  | Progressive grazing |  | Wyoming |
| *OR* |  | *OR* |  | *OR* |  | *OR* |
| Goat |  | Erosion |  | Working Lands |  | Idaho |
|  |  | *OR* |  | *OR* |  | *OR* |
|  |  | Nutrient Cycling |  | Working Landscapes |  | Utah |
|  |  | *OR* |  | *OR* |  | *OR* |
|  |  | Sequest* |  | Traditional |  | Colorado |
|  |  | ________________ |  |  |  | *OR* |
|  |  | Runoff |  |  |  | North Dakota |
|  |  | *OR* |  |  |  | *OR* |
|  |  | Water quality |  |  |  | South Dakota |
|  |  | *OR* |  |  |  | *OR* |
|  |  | Nutrient runoff |  |  |  | Nebraska |
|  |  | *OR* |  |  |  | *OR* |
|  |  | Eutrophication |  |  |  | Florida |
|  |  | ________________ |  |  |  | *OR* |
|  |  | Climate change |  |  |  | Washington |
|  |  | *OR* |  |  |  | *OR* |
|  |  | Global Warming |  |  |  | Oregon |
|  |  |  |  |  |  | *OR* |
|  |  | Trampl* |  |  |  | California |
|  |  |  |  |  |  | *OR* |
|  |  | Traditional Ecological Knowledge |  |  |  | New Mexico |
|  |  | *OR* |  |  |  | *OR* |
|  |  | Indigenous Knowledge  *OR*  Indigenous  *OR*  American Indian  *OR*  Native American |  |  |  | Arizona |

**Table S2.** Examples of the type of finding that would classify a paper into one of our six value categories.

| **Category** | **Example:** |
| --- | --- |
| Negative | Livestock are a threat to rare plants. |
| Dependent/Negative | Continuous grazing severely increases soil compaction, but there is less compaction with rotational grazing. |
| Dependent | Grazing can affect different species in different ways, particularly if other factors such as hunting or natural gas are present. In this model, elk and grass will decrease and sage grouse will increase. |
| Dependent/Positive | Grazing for heterogenous vegetation structure can help sustain bird diversity, but vegetation heights should be monitored for optimal results. |
| Positive | Native plant richness was higher in grazed plots than un-grazed plots. |
| Neutral | Grazing had no effect on Northern bobwhite habitat selection. |

**Table S3.** To understand how environmental factors play out by ecoregion we calculated representative values for each intersection based on an average of results. This table shows an example of this process for how grazing effects wildlife on the Great Plains.

| Step 1: Calculate the percent of papers in each group: | | | | | |
| --- | --- | --- | --- | --- | --- |
| Negative | Dependent/Negative | Dependent | Dependent/Positive | Positive | Neutral |
| 0.172 | 0.110 | 0.352 | 0.166 | 0.200 | .097 |
| Step 2: Multiply each category by the following corresponding values: negative=-1, dependent negative=-.5, dependent= 0, dependent/positive=.5, positive=1 (do not include neutral) | | | | | |
| Negative | Dependent/Negative | Dependent | Dependent/Positive | Positive | |
| -0.172 | -0.055 | 0 | 0.083 | 0.200 | |
| Step 3: Find the sum of the categories. The maximum value would be 1 if all papers were positive. The minimum value would be -1 if all papers were negative. | | | | | |
| Total: 0.056 | | | | | |

**Figure S1.** This figure shows the same information as Figure 2b, with the added dimension of sample size. Bubble size is representative of the number of papers in that respective category. Results are grouped by ecoregion and colors correspond to the environmental question represented. Values are explained in Table S3 with 1 denoting all positive papers and -1 all negative.

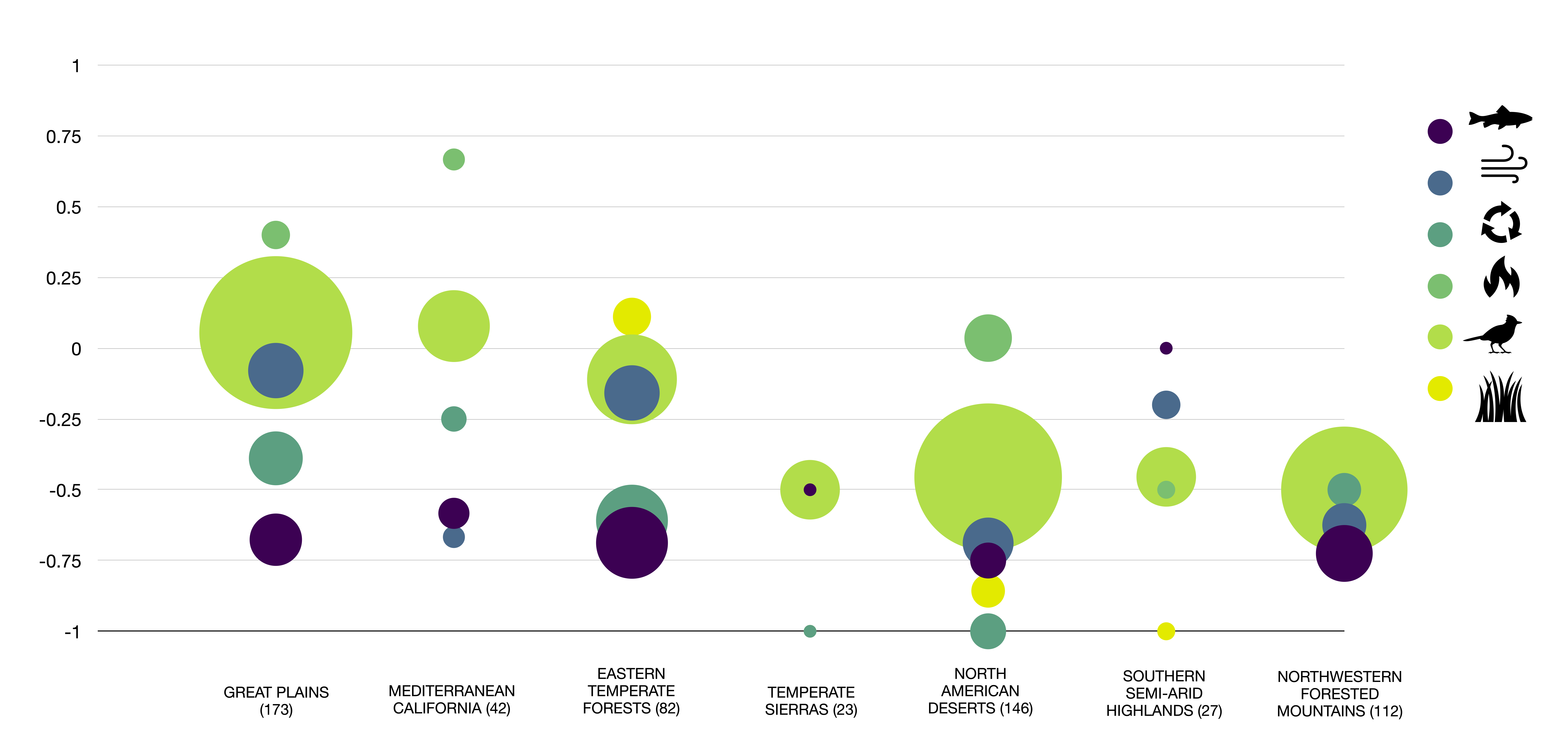
